# The gene signatures of human alpha cells in types 1 and 2 diabetes indicate disease-specific pathways of alpha cell dysfunction

**DOI:** 10.1101/2022.02.22.481528

**Authors:** Emanuele Bosi, Piero Marchetti, Guy A. Rutter, Decio L. Eizirik

## Abstract

Glucagon secretion is perturbed in both type 1 and type 2 diabetes (T1D, T2D) the pathophysiological changes at the level of individual pancreatic alpha cells are still largely obscure. Using recently-curated single-cell RNA data from human donors with either T1D or T2D and appropriate controls, we leveraged alpha cell transcriptomic alterations consistent with both common and discrete pathways. Firstly, altered expression of genes associated with alpha cell identity (*ARX, MAFB*) was common to both diseases. In contrast, increased expression of cytokine-regulated genes and genes involved in glucagon biosynthesis and processing were apparent in T1D, whereas mitochondrial genes associated with reactive oxygen species generation (*COX7B, NQO2*) were dysregulated in alpha cells from T2D patients. Conversely, T1D alpha cells displayed alterations in genes associated with autoimmune-induced ER stress (*ERLEC1, HSP90*) whilst those from T2D patients showed changes in glycolytic and citrate cycle genes (*LDH, PDHB, PDK4*) which were unaffected in T1D. These findings suggest that despite some similarities related to loss-of-function, the alterations of alpha cells present important disease-specific signatures, suggesting that they are secondary to the main pathogenic events characteristic to each disease, namely immune-mediated-or metabolic-mediated-stress in respectively T1D and T2D.

## Introduction

Both types 1 (T1D) and 2 diabetes (T2D) are characterised by varying degrees of pancreatic beta cell failure (Eizirik, Pasquali, and Cnop 2020; Marchetti et al. 2020). This is paralleled by dysfunction of alpha cells, which in T1D may contribute to insulin-induced hypoglycaemia and in T2D, at least at the initial phases of the disease, to hyperglycemia (Brissova et al. 2018; Gromada, Chabosseau, and Rutter 2018).

Alpha and beta cells are intermingled in human pancreatic islets (Bosco et al. 2010) and there is a crosstalk between these cells that regulates at least in part their function (Campbell and Newgard 2021). It is thus conceivable that the reduced functional beta cell mass in T1D and T2D impacts alpha cells and contributes to their dysfunction in each disease. Alternatively, it may be that mechanisms inherent to each disease, i.e. a predominance of autoimmunity and consequent islet inflammation in T1D as compared to severe metabolic stress in T2D (Eizirik, Pasquali, and Cnop 2020), directly impair the alpha cells.

Differentiated cells trigger diverse adaptive responses that are determined by the stress to which they are exposed. For instance, beta cells exposed to pro-inflammatory cytokines trigger branches of the unfolded protein response that are different from the ones triggered in response to the metabolic stressor palmitate, and the global gene signatures of islets obtained from patients affected by T1D or T2D are markedly different (Eizirik, Pasquali, and Cnop 2020). The cellular responses to diverse stresses may leave gene expression footprints – particularly in long-lived cells such as human alpha and beta cells – that can be detected by RNA sequencing. Examination of these footprints may enable to differentiate the principal cause(s) of the alpha cell stress present in T1D and T2D. If the leading cause of alpha cell stress is the relative or absolute loss of neighbouring beta cells, and the deficiency of insulin leading to hyperglycemia, we may expect to find similar gene signatures on alpha cells from T1D or T2D patients; on the other hand, if the stress is disease-specific then alpha cells should show different signatures in each case, for instance immune-induced stress in T1D and more metabolic changes T2D.

To test these hypotheses we presently used recently-curated human islet single-cell transcriptomic data from control donors or individuals affected by either T1D or T2D that are publicly available (Kaestner et al. 2019). The results indicate similar patterns, but also major divergences between the gene expression signatures present in alpha cells from T1D or T2D patients, arguing in favour of disease-specific mechanisms leading to alpha cell dysfunction in each case.

## Materials and Methods

### Analysis of single-cell data and integration with the other datasets

Fastq files and the corresponding metadata were downloaded from the database of the Human Pancreas Analysis Program (HPAP) (https://hpap.pmacs.upenn.edu). Reads were aligned using STAR 2.7.3a (Dobin et al. 2013) against the human reference genome GRCh37 (Ensembl 87 annotation) with different parameters according to the technology used for library production. For the reads from libraries prepared with Fluidigm 800 cell IFC, the mapping parameters were “--quantMode TranscriptomeSAM GeneCounts, --outSAMmultNmax -1”; for the other samples, prepared using 10X with chemistry Single Cell 3’ v2 or v3, the parameters used were “--soloType Droplet --soloUMIfiltering MultiGeneUMI --soloCBmatchWLtype 1MM_multi_pseudocounts --outSAMmultNmax -1 --outSAMtype BAM SortedByCoordinate --quantMode TranscriptomeSAM GeneCounts”. The arguments of two parameters, “--soloCBwhitelist” and “--soloUMIlen“, were different for the 10X chemistry kits: “737K-august-2016.txt” and “10” for v2; “3M-february-2018.txt” and “12” for v3.

After the read mapping step, read count tables were analysed with ad-hoc python scripts implementing the toolbox Scanpy (Wolf, Angerer, and Theis 2018). In particular, cell-wise and gene-wise metrics were computed for quality control analyses (QC) to define excluding criteria of low-quality cells, as previously done (Bosi et al. 2020). The parameters considered for each cell were: i) the number of genes with at least one read mapped (*expressed genes*); ii) the number of reads mapped to genes (*counts*); iii) the ratio of reads mapped on mitochondrial genes (*mitochondrial fraction*). Additionally, genes with detected expression in three or fewer cells were excluded from the analyses. These values were used to exclude cells with high counts and expressed genes (likely representing multiplets), and with high mitochondrial fraction and low expressed genes (indicative of lysed cells). These variables and their covariation were considered separately for each sample, defining separate threshold values to flag cells as “low quality”.

After QC, the processed samples were concatenated and the counts normalized to a total of 10,000 for each cell, then log-transformed. The dispersion of each gene with respect to its mean value was computed in the integrated dataset to annotate highly variable genes using the homonymous Scanpy function. The resulting set of genes displaying high variability was used to perform dataset integration with Batch Balanced KNN (BBKNN) (Polański et al. 2020), similarly as done by Park and colleagues (Park et al. 2020). Briefly, BBKNN was first used with the HPAP id as batch key, followed by a Louvain clustering (Lu, Halappanavar, and Kalyanaraman 2015) on the corrected dataset. Then, the obtained clusters were used as biological covariates to perform a ridge regression on the adjusted data, followed by a further BBKNN correction. The transcriptomes of single-cells were visualized using the Uniform Manifold Approximation Projection (UMAP).

### Unsupervised cell type annotation

Single-cell clusters were identified with the Louvain modularity algorithm (Lu, Halappanavar, and Kalyanaraman 2015) as implemented in Scanpy (https://github.com/vtraag/louvain-igraph) with resolution=0.5, with further sub-clustering for groups of interest. The genes with cluster-specific expression were found with the “rank_genes_groups” function of Scanpy, curating their association with known cell types using literature information and gene expression markers reported in PanglaoDB (Franzén, Gan, and Björkegren 2019).

### Differential expression and gene set enrichment analysis

The differentially expressed genes (DEGs) in T1D and T2D as compared to the corresponding controls were identified using MAST (Finak et al. 2015) with the following mixed effect model: *Counts ∼ Diabetes + ngenes + Technology + Race + Sex + BMI + Age*, including also a random effect estimated for each individual (*Individual*). Since the estimation of such an effect may be hampered by a low number of observations, individuals with < 50 cells were excluded from the analysis. *Counts* is the matrix of raw count data, filtered of genes being expressed in less than 20% of the cells with the *filterLowExpressedGenes* function, *ngenes* is a variable encoding the number of expressed genes and *Diabetes* is a 2-level factor (Disease, ND) indicating the diabetes status of the donor. Genes with corrected p-value (Benjamini-Hochberg, FDR) lower than 0.05 and absolute fold-change (FC) greater or equal than 0.5 were considered as DEGs.

Gene set enrichment analysis (GSEA) was performed for the following datasets: i) Gene Ontology (GO), separately for Biological Process (GO BP), Molecular Function (GO MF) and Cellular Component (GO CC) (Ashburner et al. 2000; The Gene Ontology Consortium 2019); ii) KEGG (M. Kanehisa and Goto 2000; Minoru Kanehisa et al. 2019; Minoru Kanehisa 2019); iii) Reactome 2016 (Jassal et al. 2020); the Hallmark collection from mSigDB (Liberzon et al. 2015). All gene sets were obtained from mSigDB (Subramanian et al. 2005) (version 7.4) and used to perform GSEA on the hurdle distributions obtained with MAST: the “gseaAfterBoot” function has been used on 50 distributions (bootstraps) derived with the “bootVcov1” function.

### Assembling single-cell datasets from the HPAP database

The populations of T1D and T2D donors are characterized by different clinical features, with the most prominent differences being in age and body mass index (BMI), which usually are higher in T2D (American Diabetes Association 2010). For this reason, separate control groups were assembled to match (as much as possible) the clinical features of the affected donors, controlling: i) single-cell technology used to assess transcriptomes, ii) age, iii) BMI and iiii) sex. Normoglycemic donors showing positivity towards pancreatic auto-antibodies were excluded. After the selection of control donors on the basis of such criteria, the T1D dataset included 7 diabetic donors and 6 controls, whereas T2D contained 5 affected donors and 5 controls (**Figure 1A**). The features of the donors included in this study are reported in **Supplementary Table 1**.

**Figure 1.**
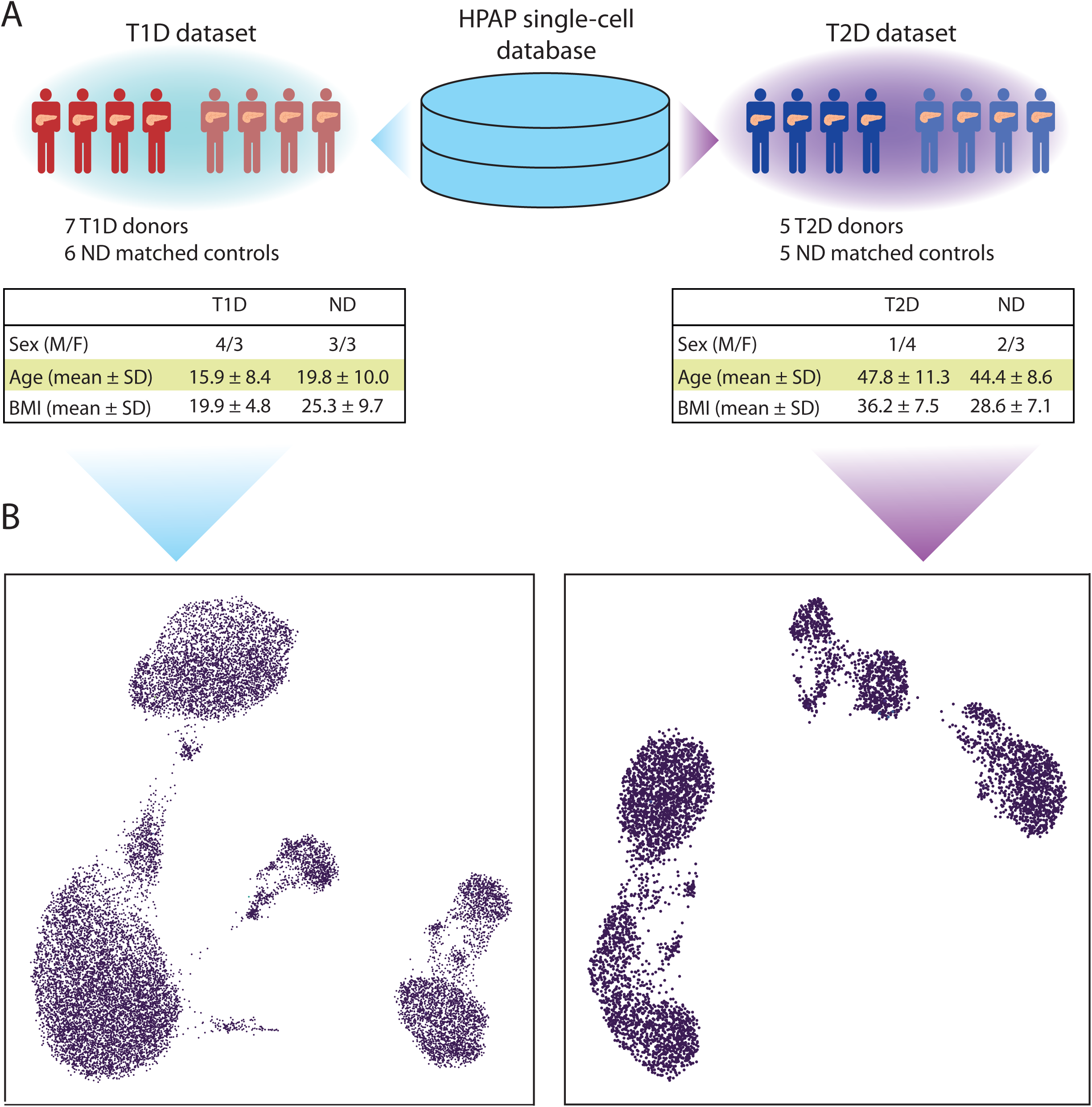
T1D and T2D single-cell transcriptomics datasets. **A:** Single-cell transcriptomic samples were obtained from the database of Human Pancreas Analysis Program (HPAP) and divided among two datasets, T1D and T2D. Samples from non-diabetic donors were assigned to a dataset according to donor and technical features in order to match those of the T1D and T2D samples. The tables report the donor features of T1D and T2D datasets. **B:** The UMAP plots provide a representation of the T1D (left) and T2D (right) datasets. Each dot corresponds to the transcriptomic profile of a single-cell projected in a two dimensional space (UMAP1, UMAP2).

Analysis of raw sequencing data and exclusion of low-quality cells and genes (see Methods) delivered a T1D dataset comprising 17,503 cells and 31,890 genes, and a T2D dataset with 5,772 cells and 32,221 genes. The QC thresholds are reported in **Supplementary Table 2**. The datasets have been corrected for technical covariates (single-cell technology and sequencing batch) and visualised in a UMAP space (**Figure 1B, Supplementary Figure 1, Supplementary Figure 2**). Clusters of single-cells were identified with the Louvain algorithm and annotated considering the expression of marker genes (**Figure 2**). The genes highly expressed in the annotated clusters were compared with the marker genes reported in PanglaoDB (Franzén, Gan, and Björkegren 2019) to confirm the cell type annotation.

**Figure 2.**
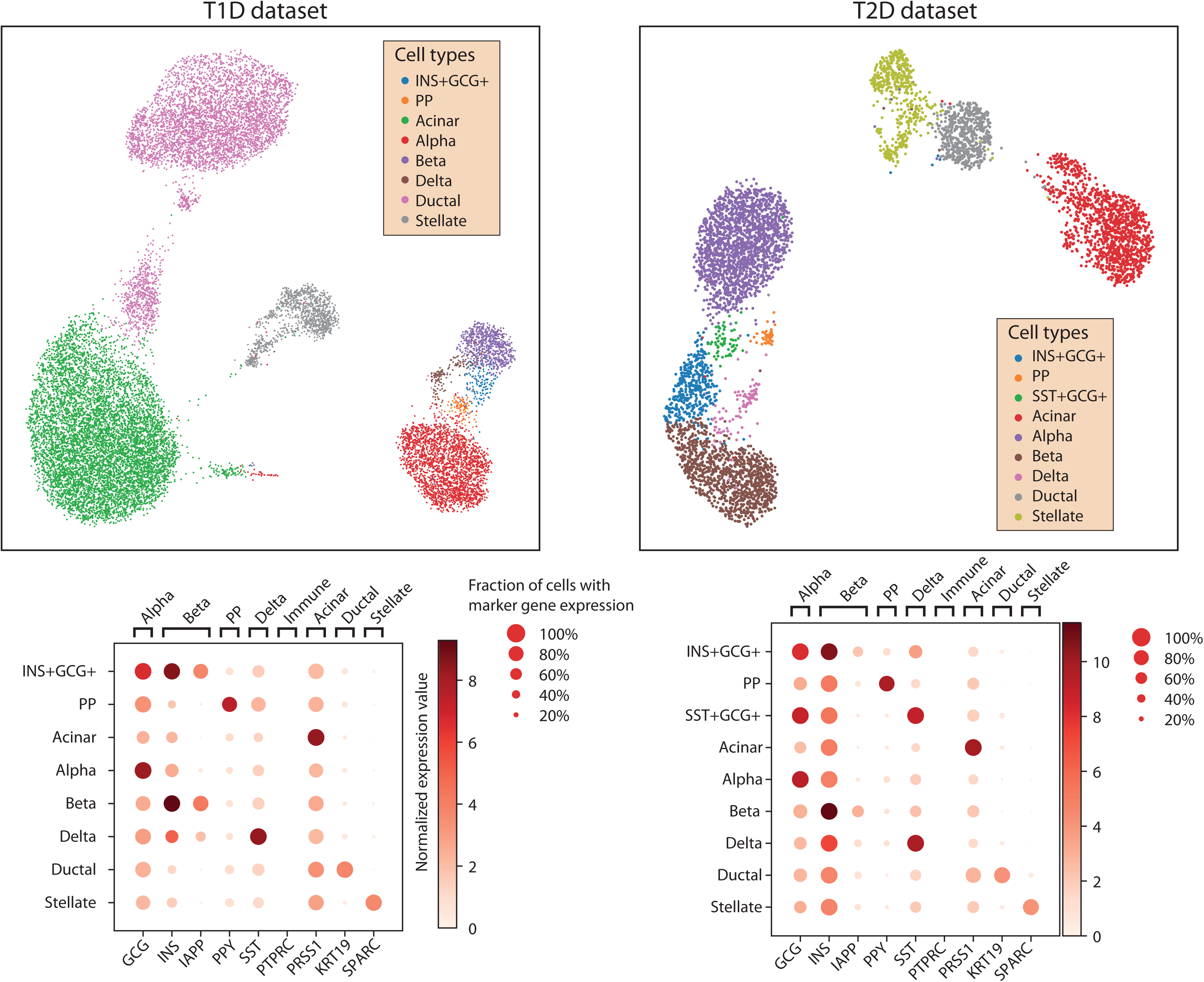
Cell type annotation of T1D and T2D datasets. The single-cells in the UMAP plots of T1D (left) and T2D (right) datasets have been clustered and annotated to a pancreatic cell type (top) based on the expression of known marker genes (bottom).

The number of cells for each cell type in each dataset are reported in **Table 1**. Alpha cells were then analysed separately in each dataset to find transcriptomic alterations potentially linked to diabetes.

**Table 1.** Cell type distributions in the T1D and T2D datasets. The table reports the number of cells assigned to each cell type in the T1D and T2D datasets

|  | INS+GCG+ | SST+GCG+ | PP | Acinar | Alpha | Beta | Delta | Ductal | Stellate |
| --- | --- | --- | --- | --- | --- | --- | --- | --- | --- |
| T1D | 140 | 0 | 97 | 7909 | 2362 | 736 | 145 | 4996 | 1118 |
| T2D | 382 | 61 | 43 | 1246 | 1757 | 1098 | 102 | 564 | 518 |

### The transcriptional signatures of alpha cells in T1D

The T1D dataset comprised 2,362 alpha cells, collectively expressing 6,265 genes (after filtering). A more detailed break-down of the number of cells at the level of individual, single-cell technology used and diabetes status is reported in **Supplementary Table 3**. After removing cells from individuals with less than 50 transcriptomes, the resulting dataset, including 2,225 alpha cells, was analysed with MAST to identify genes differentially expressed in T1D cells.

A total of 346 DEGs were identified, of which 193 were up-regulated and 153 were down-regulated versus cells from the control, normoglycemic group (**Supplementary Table 4**). A pattern observed among the DEGs was the over-expression of genes involved in Reactive Oxygen Species (ROS) response (*PRNP, NDUFA6, NDUFB4, GLRX* and *TXN*), and protein folding stability via chaperone activity (*HSP9, HSP90AA1* and *HSP90AB1*). We also observed overexpression of IL-8 and HLA-A, genes downstream of the transcription factors NF-κB and STAT1/STAT2 that are regulated by the pro-inflammatory cytokines IL-1β and types I and II interferons (IFNs) and participate in the immune system-islet cell dialogue present in T1D (Eizirik, Pasquali, and Cnop 2020). Importantly, these genes were not overexpressed in alpha cells from patients affected by T2D (see below). The over-expression of DDIT3 (also known as CHOP), a key mediator of ER stress induced-beta cell death (Eizirik, Cardozo, and Cnop 2008), fits with this scenario. Another trend of pathophysiological relevance was the significant down-regulation of genes implicated in endocrine function, namely PCSK2, CHGA, SRP14 and PAK3. These genes contribute to proglucagon peptide maturation and eventual exocytosis, and their decreased expression could contribute to reduced glucagon secretion from alpha cells in human T1D (Gerich et al. 1973). Thus, in PCSK2 gene-null mice lacking the prohormone convertase in alpha cells, proglucagon maturation to the mature hormone is blocked (Furuta et al. 2001). Of interest, there was also down-regulation (−2.5 fold change) of PCSK1N, a specific inhibitor of PCSK1.

A comprehensive investigation focusing on the functional signatures enriched in T1D was performed with an enrichment test as implemented in MAST, using six different collections (mSIGDB, Reactome, KEGG, GO-MF, GO-BP, GO-CC) encompassing broad (i.e. mSIGDB, KEGG) or very specific (i.e. GO) functional terms (**Supplementary Table 5)**. Collectively, there were a total of 1,539 significantly enriched terms, with 1,159 positively- and 380 negatively enriched. Of these, 1,207 were GO terms (159 MF, 896 BP, 152 CC), whereas mSIGDB, KEGG and Reactome presented 22, 49 and 261 enriched terms. Some of these terms were consistent with the patterns observed with the results of the differential expression analysis, in that functional categories related to ROS and unfolded protein response (UPR) and exposure to immune mediators were consistently enriched in the different datasets, thus reinforcing the view that T1D alpha cells endure higher levels of these stresses. To better highlight the most important signatures of T1D alpha cells, the enrichment results were ranked according to the obtained significance. The top three KEGG categories positively enriched were related to immunity (ALLOGRAFT REJECTION, AUTOIMMUNE THYROID DISEASE, ANTIGEN PROCESSING AND PRESENTATION), consistently with the inflammation associated with islets in T1D (Eizirik, Pasquali, and Cnop 2020). The most positively enriched KEGG terms also included RIBOSOME and OXIDATIVE PHOSPHORYLATION, both associated with increased ROS production. Mitochondrial activity is a major source of ROS, whereas oxidative stress impairs protein biosynthesis and modifies rRNA (Shcherbik and Pestov 2019), requiring an increased turnover of these molecules by activating mechanisms to repair or recycle damaged molecules. Pathways associated with immunity and ROS production/response were among the most positively enriched pathways in all databases. In Reactome we found, related to immunity, a positive enrichment of interferon gamma signalling, endosomal/vacuolar pathway of antigen presentation and trafficking/processing of endosomal toll-like receptor (TLR). Related to ROS there is an increase of mitochondrial fatty acid oxidation, metallothionein ROS scavenging, and actin folding. In mSIGDB hallmark the top ten pathways included allograft rejection, oxidative phosphorylation, apoptosis, interferon gamma response, unfolded protein response and reactive oxygen species. The top three GO biological processes were negative regulation of cell killing, negative regulation of leukocyte mediated cytotoxicity and positive regulation of T-cell mediated cytotoxicity. The other highly enriched pathways include positive regulation of protein depolymerization, positive regulation of actin filament depolymerization and negative regulation of IRE1-mediated unfolded protein response.

### The transcriptional signatures of alpha cells in T2D

After filtering out low quality cells, in the T2D dataset there were 1,757 alpha cells expressing a total of 6,872 genes. Following the approach used for the T1D dataset, we detailed the number of alpha cells at the levels of individual, single-cell technology and diabetes status (**Supplementary Table 3**). Two donors had a low number of cells and were excluded from the differential expression analysis; the remaining cells (1,718) were analysed with MAST.

There were 466 genes differentially expressed in T2D versus control alpha cells, 238 up-regulated and 228 down-regulated (**Supplementary Table 6**). Among the over-expressed DEGs, there were activators of stress and apoptosis mediated by p53, i.e. *CCNK, JUN, BTG1, GADD45A, GADD45B, XPC, CDKN2AIP, DDIT3, TXNIP*, and *ATF3*. Of these, *XPC, GADD45A, GADD45B* and *CDKN2AIP* respond to DNA oxidative damage (Wang et al. 2012;

Salvador, Brown-Clay, and Fornace 2013; Hasan et al. 2009), while *DDIT3* is activated by endoplasmic reticulum (ER) stress (Yamaguchi and Wang 2004; Ohoka et al. 2005; Eizirik, Cardozo, and Cnop 2008). *TXNIP* and *ATF3* are instead modulated by glucose concentration and play a role in both apoptosis and hormone secretion. TXNIP controls a pathway (miR-204/MafA/insulin) that reduces insulin expression (Xu et al. 2013), whereas ATF3 increases the expression of proglucagon, as well as regulating expression of genes related to apoptosis (such as *GADD45, BNIP3* and *NOXA*).

The over-expression of such genes, and others related to ROS (*SBNO2, EGLN2, MBP*) and the unfolded protein response (UPR; *EEF2, ATF4, HERPUD1*) suggests that alpha cells in T2D are subject to oxidative stress, a hallmark of toxicity induced by elevate glucose levels (Kawahito, Kitahata, and Oshita 2009; Gromada, Chabosseau, and Rutter 2018). The down-regulation of genes involved in pyruvate metabolism (*LDHA, PDHB, PDK4*) and oxidative phosphorylation (*COX7B, NQO2, SUCLA2, UQCR10, SLC25A4*) is also consistent with oxidative stress pathways, in that these may affect ROS production by mitochondria.

Of functional relevance to hormone biosynthesis, the genes encoding prohormone convertases *PCSK1* and *PCSK2* were both down-regulated, as well as SCGN and SYT13, involved in hormone storage and exocytosis, respectively (Yang et al. 2016; Andersson et al. 2012), and the transcription factor MAFB, crucial for preproglucagon gene expression and the production and release of the hormone (Katoh et al. 2018).

As for the T1D dataset, a gene set enrichment analysis was performed to identify functions and pathways significantly enriched in the T2D alpha cells. Overall, 1,973 significantly enriched terms were identified, of which 494 positively and 1,479 negatively enriched (**Supplementary Table 7**). Most of these (1,552) were Gene Ontology (GO) terms (207 MF, 1158 BP, 187 CC), while the enriched terms in mSIGDB, KEGG and Reactome were 12, 99 and 310, respectively.

The 6 mSIGDB pathways positively enriched (“INTERFERON GAMMA RESPONSE”, “INTERFERON ALPHA RESPONSE”, “P53 PATHWAY”, “ALLOGRAFT REJECTION”, “TNFA SIGNALING VIA NFKB”, “APOPTOSIS”) are associated to inflammation, whereas the negatively enriched terms are mostly involved in the energetic metabolism (“OXIDATIVE PHOSPHORYLATION”, “ADIPOGENESIS”, “GLYCOLYSIS”). This trend is consistent in the other sets: for instance, in KEGG the terms such as “P53 SIGNALING PATHWAY” and “TOLL LIKE RECEPTOR SIGNALING PATHWAY” are positively enriched, while negatively enriched ones included metabolic terms such as “PYRUVATE METABOLISM”, “KEGG STARCH AND SUCROSE METABOLISM” and “CITRATE CYCLE TCA CYCLE”. Similarly, among the Reactome positively enriched pathways there are “INTERFERON GAMMA SIGNALING”, “INFLAMMASOMES” and “ANTIGEN PRESENTATION FOLDING ASSEMBLY AND PEPTIDE LOADING OF CLASS I MHC”, while the negatively enriched ones include “REGULATION OF PYRUVATE DEHYDROGENASE PDH COMPLEX”, “THE CITRIC ACID TCA CYCLE AND RESPIRATORY ELECTRON TRANSPORT” and “GLUCOSE METABOLISM”. Of interest, we report a number of negatively enriched terms specifically linked to alpha cell function: “GLUCAGON SIGNALING IN METABOLIC REGULATION”, “PKA ACTIVATION IN GLUCAGON SIGNALLING”, “GLUCAGON LIKE PEPTIDE 1 GLP1 REGULATES INSULIN SECRETION” and “GLUCAGON TYPE LIGAND RECEPTORS”, among the Reactome terms; “CELLULAR RESPONSE TO GLUCAGON STIMULUS” and “RESPONSE TO GLUCAGON” among the GO-BP terms.

### Contrasting the alpha cell signatures of T1D and T2D

By analysing the changes in gene expression in alpha cells separately in T1D and T2D the corresponding transcriptomic signatures were identified and described, revealing a number of similarities, such as upregulated DEGs involved in the stress response or the downregulation of genes relevant for the secretory function. To quantify more precisely the extent at which the two series overlap and to underscore their differences, a systematic comparison of the obtained results was undertaken next.

In T1D and T2D there were a total of 770 DEGs (**Supplementary Table 8**). Of these, 42 were shared, while DEGs present only in T1D and T2D were 304 and 424, respectively. Considering only up-regulated genes, there were 420 DEGs, of which 11 were shared, 182 were T1D-specific and 227 were T2D-specific (**Figure 3**). The shared up-regulated genes are *EIF4A2, DDIT3, RRAGD, RPL26, SNHG6, C9orf16, KIF1A, ZNF706, SERTAD1, RSL24D1* and *HNRNPF*. For the down-regulated genes, from a total of 368 DEGs there were 13 shared, 140 T1D-specific and 215 T2D-specific. The down-regulated DEGs in common are *C4orf48, KRT10, C1QBP, PCSK2, REG1B, PEG10, PAM, CTRB2, PRSS3P1, CTNND2, CTRB1, ARL3* and *ATXN10*. The overlap of DEGs regulated in different directions in T1D and T2D was assessed as well. There were 10 genes in common between the T1D up-regulated and the T2D down-regulated DEGs (*STXBP2, CSTB, ATP5B, SERF2, RPN2, GNG4, ATP5I, CELA3A, CD99* and *POLR2K*), while the shared genes between the T1D down-regulated and the T2D up-regulated were 8 (*IER2, JUND, TXNIP, EIF5A, INO80E, C19orf43, TRAK1* and *TBL1XR1*). Finally, since DEGs were identified based both on significance and fold-change thresholds, there are several genes that have a significant change but with fold-change below threshold. Although these genes have not been reported as DEGs, some could provide relevant information. For instance, genes related to alpha cell identity (notably the transcription factors *ARX* and *MAFB*) were significantly down-regulated in both T1D and T2D, and, although their fold-change fell below a 0.5 threshold, this provides evidence that both diseases are associated with alteration of alpha cell identity and function.

**Figure 3.**
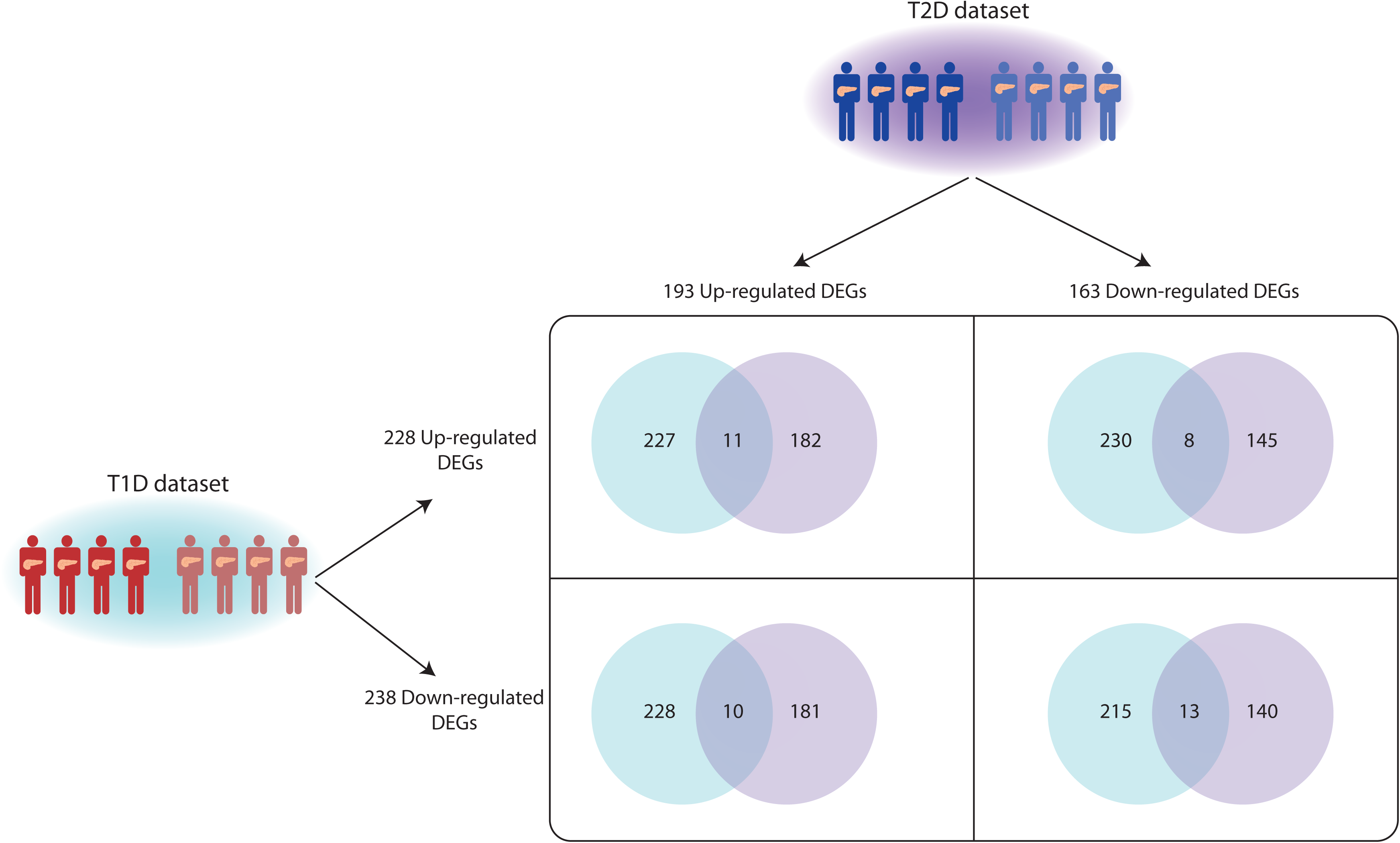
Comparison of differentially expressed genes in T1D and T2D alpha cells. Alpha cells from T1D (bottom left) and T2D (top right) datasets were separately analysed contrasting the expression signatures of cells from affected and ND individuals. The differentially expressed genes (DEGs) were compared to identify shared and dataset-specific DEGs.

**Figure 4.**
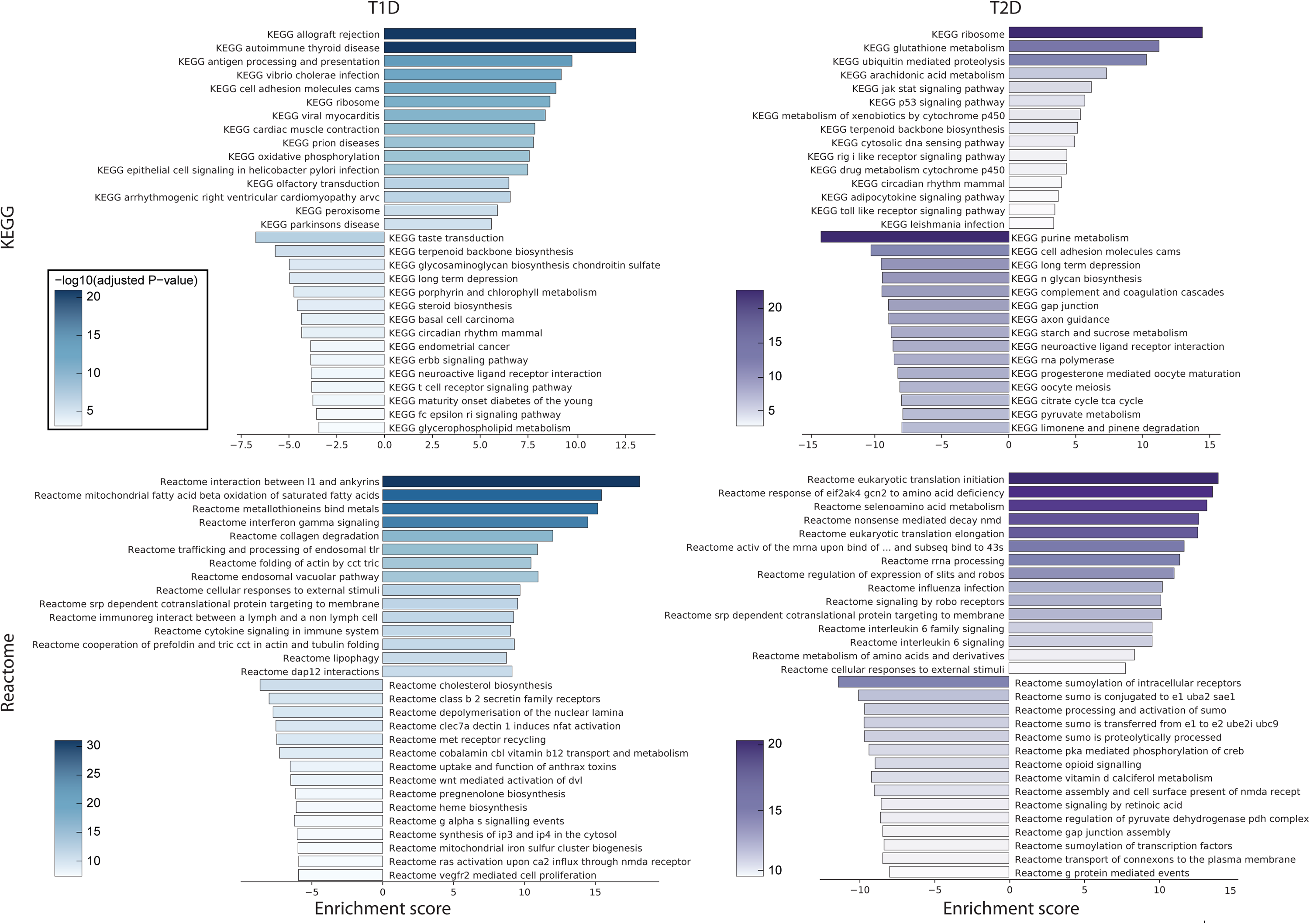
Comparison of most significantly enriched pathways in T1D and T2D. Gene-set enrichment analysis was performed for T1D (left) and T2D (right) datasets. The barplots report the normalized enrichment score (bar length) and the significance (bar color) of the most significant 15 positively and negatively enriched pathways from KEGG (top) and Reactome (bottom).

Examining the extent of the overlap between GSEA results, from a total of 2,974 terms significantly enriched either in T1D or T2D (**Supplementary Table 9**), 506 were shared, 1,033 were unique to T1D and 1,435 to T2D. Considering only the positively enriched terms (in T1D, T2D or both), there were 1,475 terms, 171 of which in common; the negatively enriched ones were 1,781 in total, with 53 shared. There were 282 terms enriched in different directions: 261 of these were positively enriched in T1D and negatively in T2D, whereas 21 were shared between positively enriched in T2D and negatively in T1D. A breakdown of the overlapping terms in T1D and T2D (in different directions, and their combinations) across the different datasets used for GSEA is reported in **Supplementary Table 9**.

Although the signatures retrieved for T1D and T2D indicate a partial overlap of stress response genes, we found some remarkable differences between the two series. In T1D, genes linked with ER stress and UPR are either significantly up-regulated (*ERLEC1, HSP90*) or with a significant change that is below the fold-change threshold we used (*ATF6*, ER mannosidase I, *TRAM1*), an effect that was not observed in T2D (**Supplementary Figure 3**).

Another major difference between T1D and T2D involved metabolism i.e. glycolysis, citrate cycle and oxidative phosphorylation (**Supplementary Figure 4**). Notably, in T1D we observed little changes in expression of genes involved glycolysis and citrate cycle, but increased levels of those contributing to oxidative phosphorylation, whilst these pathways were repressed in T2D: most of the genes involved in glycolysis were down-regulated, four with high significance (*PKM, ENO1, ALDH2, TPI1*) and three with high significance and fold-change (*LDHA, ALDH9A1* and *PDHB*); strikingly, most of the citrate cycle genes were down-regulated in alpha cells from T2D donors, with six genes displaying a significant change; among the complexes involved in oxidative phosphorylation, genes contributing in complex III were the most negatively affected, with almost all genes being under-expressed in diseased alpha cells.

## Discussion

Analysis of single-cell data sets provides a unique opportunity to better understand the changes at the level of the alpha cell which drive dysregulated glucagon secretion in T1D and T2D. In the present work we used publicly-available single-cell transcriptomic data from human islets to test whether the dysfunction of alpha cells in T1D and T2D had common bases or if the transcriptional signatures are more disease-specific. This revealed both shared features (i.e. down-regulation of genes related to alpha cell identity and function, and the up-regulation of stress response mechanisms and inflammation signatures) and, more importantly, disease-specific stress pathways. In T1D, several of the alpha cell alterations were traced back to ER stress, a process associated with chronic inflammation and autoimmune diseases. In T2D, up-regulation of ROS defense mechanisms (*SBNO2, EGLN2, MBP*) was accompanied by modification of the central metabolism, with the repression of genes involved in glycolysis (*LDHA, PDHB, PDK4*), citrate cycle and mitochondrial respiration (*COX7B, NQO2, SUCLA2, UQCR10, SLC25A4*). We note that some of these gene expression changes in T2D alpha cells are akin to findings reported in a recent study using a separate data set (Dai et al. 2022) which highlighted alteration of mitochondrial respiratory complex genes in T2D and inhibitory role of H_2_O_2_ in glucagon secretion. However, no data were provided in the latter report on alpha cells from patients with T1D, nor was any comparison made of changes in the two disease types.

We speculate that in T2D down-regulation of glucose catabolism is adaptive to ROS stress, not only by leading to a decrease of radicals from oxidative phosphorylation but also by redirecting glucose flux to Pentose Phosphate Pathway, increasing NADPH to improve ROS scavenging (Mullarky and Cantley 2015). However, the relationship here is complex since lowered LDHA (favouring pyruvate flux into mitochondria) and PDK4 (favouring PDH dephosphorylation and activation) would seem likely to oppose the impact of lowered PDHB activate (decreasing PDH-E1 levels and conversion of pyruvate into acetyl-CoA) (**Supplementary Figure 3**). Measurements of the corresponding gene products at the protein level will be important to substantiate these mRNA-based findings (**Supplementary Figure 3**). Of note, glucose oxidation is lowered in islets from T2D subjects (Del Guerra et al. 2005) though the relative contribution of changes in beta and alpha cells is not established. We also report that, among the respiratory chain genes, the most negatively down-regulated were those involved in mitochondrial complex III, a major producer of ROS with implications into cellular transduction (Bleier and Dröse 2013). Additionally, and complementing the above mechanism, a decreased ability of mitochondria to synthesise ATP in response to elevated glucose concentrations (Ravier and Rutter 2005) may contribute to the failure of the alpha cell in T2D to efficiently shut down glucagon secretion as glucose concentrations rise, consistent with other recent findings (Knudsen et al. 2019).

In T1D we observed signatures of ER stress, with most of the genes involved in ER protein processing being up-regulated (**Supplementary Figure 4**). Considering that in T1D we did not observe repression of the central metabolism (**Supplementary Figure 3**), the partial overlap of ROS response between T1D and T2D could be due to the fact that ER stress increases the ROS cellular levels (Zeeshan et al. 2016). Related to ROS production, oxidative phosphorylation is enhanced in T1D, a striking difference with respect to T2D alpha cells in which the pathway is repressed. Finally, alpha cells in T1D display signatures of inflammation, including markers of cytokine exposure, with multiple related pathways among the most significantly enriched.

Besides a stress response, we reported for T1D the down-regulation of a number of genes involved with hormone secretion, with a likely relevance in the pathophysiological context. The apparent repression of PCSK2 transcription is consistent with reduced glucagon production, as well as the decreased expression of PCSK1N, a repressor of PCSK1. As this gene is involved with glucagon-like peptide 1 (GLP-1) production from glucagon (Rouillé et al. 1997), the pattern we observed would be consistent with increased endogenous production of GLP-1 and lower glucagon maturation. Also, the down-regulation of CHGA, encoding chromogranin A, implies impaired hormone secretion with a role in secretory granule biogenesis (Kim et al. 2001).

In conclusion, despite their similarities, the alterations of alpha cells present important disease-specific signatures, suggesting that they are, at least in part, secondary to the main pathogenic events characteristic to each disease, namely immune-mediated- or metabolic-mediated-stress, respectively, in T1D and T2D. Nevertheless, we note that our findings are derived only from single-cell transcriptomics, and are therefore subject to caveats over sensitivity and other limitations associated with this approach (Mawla and Huising 2019). Future studies will be required to dissect the molecular mechanisms involved in the observed changes, their relevance for disease pathogenesis and the potential for targeted therapeutic interventions.

## Acknowledgements

This work was supported by non-profit organisations and public bodies for funding of scientific research conducted within the European Union: the Innovative Medicines Initiative 2 Joint Undertaking, RHAPSODY [115881 to EB, DLE, GAR, PM], INNODIA [115797 to EB, DLE, PM] and INNODIA HARVEST [945268 to DLE, PM] -this Joint Undertaking receives support from the Union’s Horizon 2020 research and innovation programme, “EFPIA”, “JDRF” and “The Leona M. and Harry B. Helmsley Charitable Trust” (INNODIA, INNODIA HARVEST), the “EFPIA” and the Swiss State Secretariat for Education, Research and Innovation under contract number 16.0097 (RHAPSODY); the European Union’s Horizon 2020 research and innovation programme, project T2DSystems [667191 to EB, DLE, PM]; the Walloon Region through the FRFS-WELBIO Fund for Strategic Fundamental Research [grant numbers CR-2015A-06s, CR-2019C-04 to DLE]; the Welbio-Fonds National de la Recherche Scientifique, Belgium and Dutch Diabetes Fonds, Holland [2018.10.002 to DLE]; the Brussels Capital Region-Innoviris project Diatype [2017-PFS-24 to DLE]. GAR was supported by a Wellcome Trust Investigator Award (212625/Z/18/Z), MRC Programme grant (MR/R022259/1) and a start-up grant from the CR-CHUM, Université de Montréal.

This manuscript used data acquired from the Human Pancreas Analysis Program (HPAP-RRID:SCR_016202) Database (https://hpap.pmacs.upenn.edu), a Human Islet Research Network (RRID:SCR_014393) consortium (UC4-DK-112217, U01-DK-123594, UC4-DK-112232, and U01-DK-123716).

## Figure legends

Supplementary Figure 1

**UMAP representation of T1D with confounders**

The single-cells from the T1D dataset are reported with cells color-labeled according to the single-cell technology used for sample preparation (left) and the HPAP ID of the donor (right).

Supplementary Figure 2

**UMAP representation of T2D with confounders**

The single-cells from the T2D dataset are reported with cells color-labeled according to the single-cell technology used for sample preparation (left) and the HPAP ID of the donor (right).

Supplementary Figure 3

**Expression of genes involved in Glycolysis in T1D and T2D datasets**

The KEGG map of the Glycolysis pathway (hsa00010) is reported with the enzymes colored according to the fold-change of the corresponding genes in T1D (left) and T2D (right) datasets.

Supplementary Figure 4

**Expression of genes involved in Unfolded Protein Response in T1D and T2D datasets** The KEGG map of the Unfolded Protein Response pathway (hsa04141) is reported with the enzymes colored according to the fold-change of the corresponding genes in T1D (left) and T2D (right) datasets.

Supplementary Table 1

**Features of donors in T1D and T2D datasets**

The sheet “Donor Features” reports the information of each donor included in the study. This table was used to create a pivot table (sheet “Features of T1D and T2D datasets”), with descriptive statistics (sex, age, BMI, single-cell technology) for T1D and T2D datasets.

Supplementary Table 2

**Thresholds used to filter low-quality single-cell transcriptomes**

The table reports the filters used for each single-cell sample to flag low-quality cells.

Supplementary Table 3

**Alpha cell count table**

The table reports the number of high-quality alpha cells identified in each sample, together with features associated with each donor, that are Dataset (T1D vs T2D), Diabetes (T1D, T2D, ND) and Single-cell Technology (Fluidigm, 10X v.2, 10X v.3).

Supplementary Table 4

**Alpha cell genes differentially expressed in T1D**

The table reports the results of the differential expression analysis of alpha cells in the T1D dataset. The reported DEGs are sorted according to their differential expression adjusted significance (FDR).

Supplementary Table 5

**Results of gene set enrichment analysis in T1D**

The table reports the results of gene set enrichment analysis performed using different datasets (mSIGDB, KEGG, Reactome, Gene Ontology - Molecular Function, Biological Process and Cellular Component). In particular, the table includes for each enriched term the results of MAST enrichment analysis, with the effect and significance of each component of the statistical model; the terms are sorted according to their enrichment direction (1=positively enriched in T1D, -1=negatively enriched in T1D) and significance (“combined_adj”). The results are reported for each dataset in a separate sheet, whereas the aggregated results for all the datasets are included in the “All” sheet.

Supplementary Table 6

**Alpha cell genes differentially expressed in T2D**

The table reports the results of the differential expression analysis of alpha cells in the T2D dataset. The reported DEGs are sorted according to their differential expression adjusted significance (FDR).

Supplementary Table 7

**Results of gene set enrichment analysis in T2D**

The table reports the results of gene set enrichment analysis performed using different datasets (mSIGDB, KEGG, Reactome, Gene Ontology - Molecular Function, Biological Process and Cellular Component). In particular, the table includes for each enriched term the results of MAST enrichment analysis, with the effect and significance of each component of the statistical model; the terms are sorted according to their enrichment direction (1=positively enriched in T2D, -1=negatively enriched in T2D) and significance (“combined_adj”). The results are reported for each dataset in a separate sheet, whereas the aggregated results for all the datasets are included in the “All” sheet.

Supplementary Table 8

**Differentially expressed genes shared between T1D and T2D**

Partitioning DEGs in T1D and T2D according to their direction (up- or down-regulated) allowed the comparison between up-regulated genes in T1D and in T2D, down-regulated genes in T1D and T2D, up-regulated in T1D and down-regulated in T2D, and down-regulated in T1D and up-regulated in T2D. The results of such comparisons are reported in each sheet, with a gene classified as “Shared” if it is differentially expressed in both conditions, “unique to T1D” or “unique to T2D” if it is differentially expressed only in one condition.

Supplementary Table 9

**Comparison of GSEA enriched terms between T1D and T2D**

Partitioning enriched functional terms in T1D and T2D according to their direction (up- or down-regulated) allowed the comparison between up-regulated genes in T1D and in T2D, down-regulated genes in T1D and T2D, up-regulated in T1D and down-regulated in T2D, and down-regulated in T1D and up-regulated in T2D. The results of such comparisons are reported in each sheet, with a gene classified as “Shared” if it is differentially expressed in both conditions, “unique to T1D” or “unique to T2D” if it is differentially expressed only in one condition.

