## Supplementary figures and images for "The gene signatures of human alpha cells in types 1 and 2 diabetes indicate disease-specific pathways of alpha cell dysfunction"

### Supplementary Figure 1

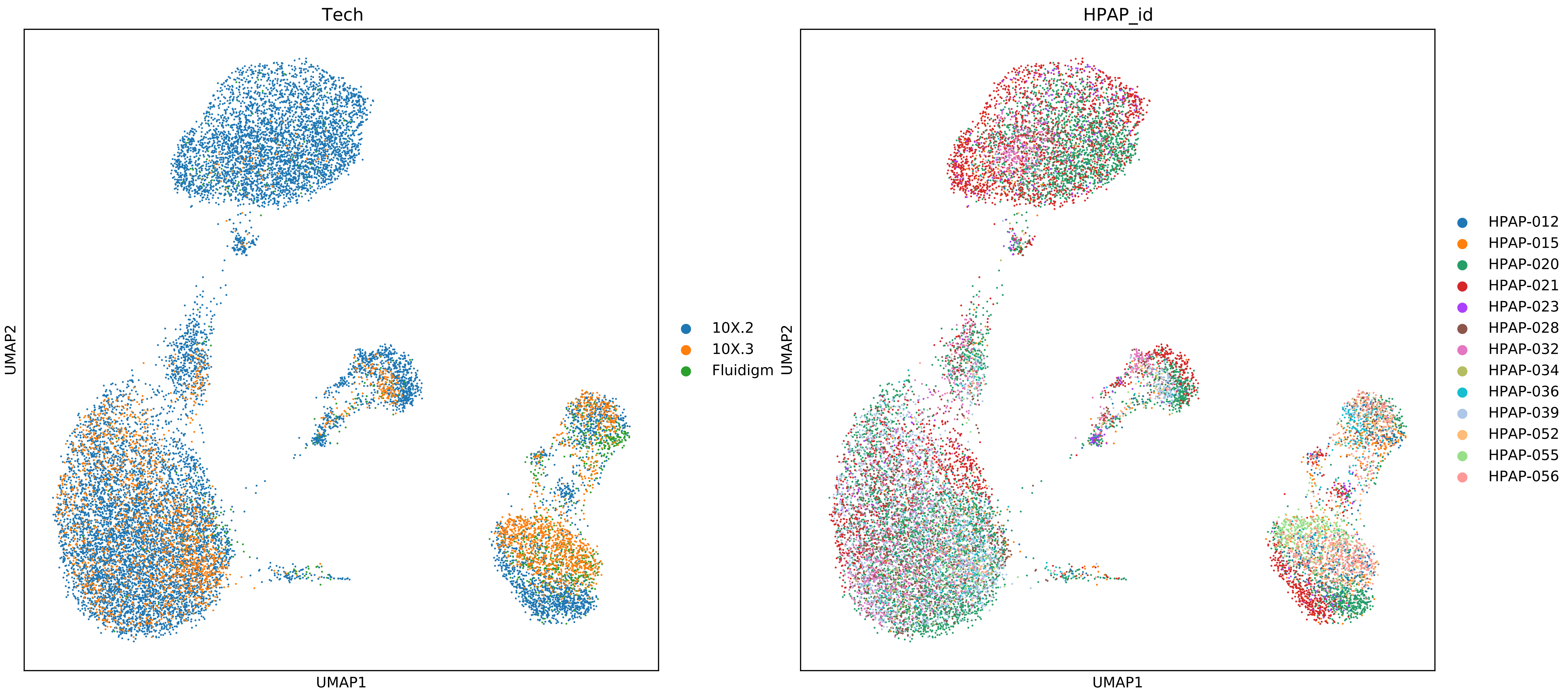

### Supplementary Figure 2

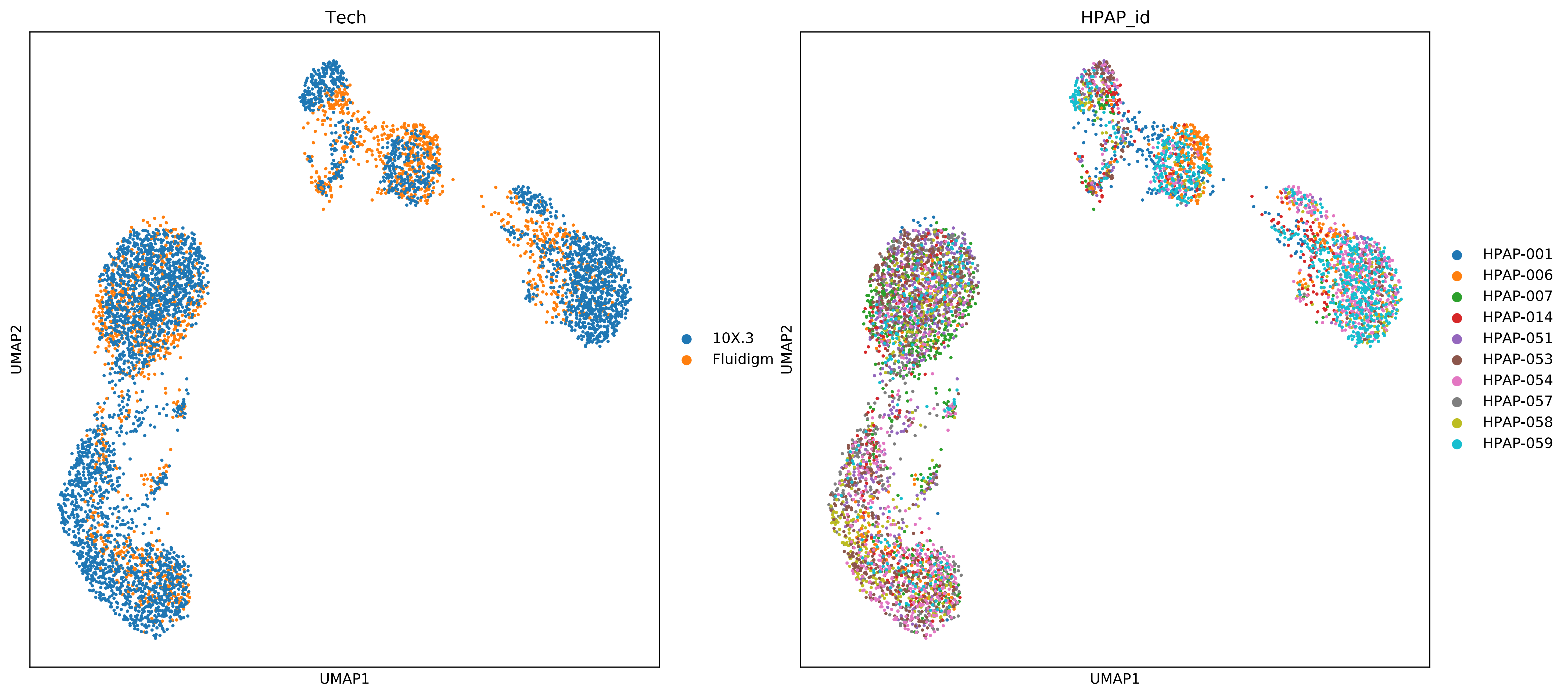

### Supplementary Figure 3

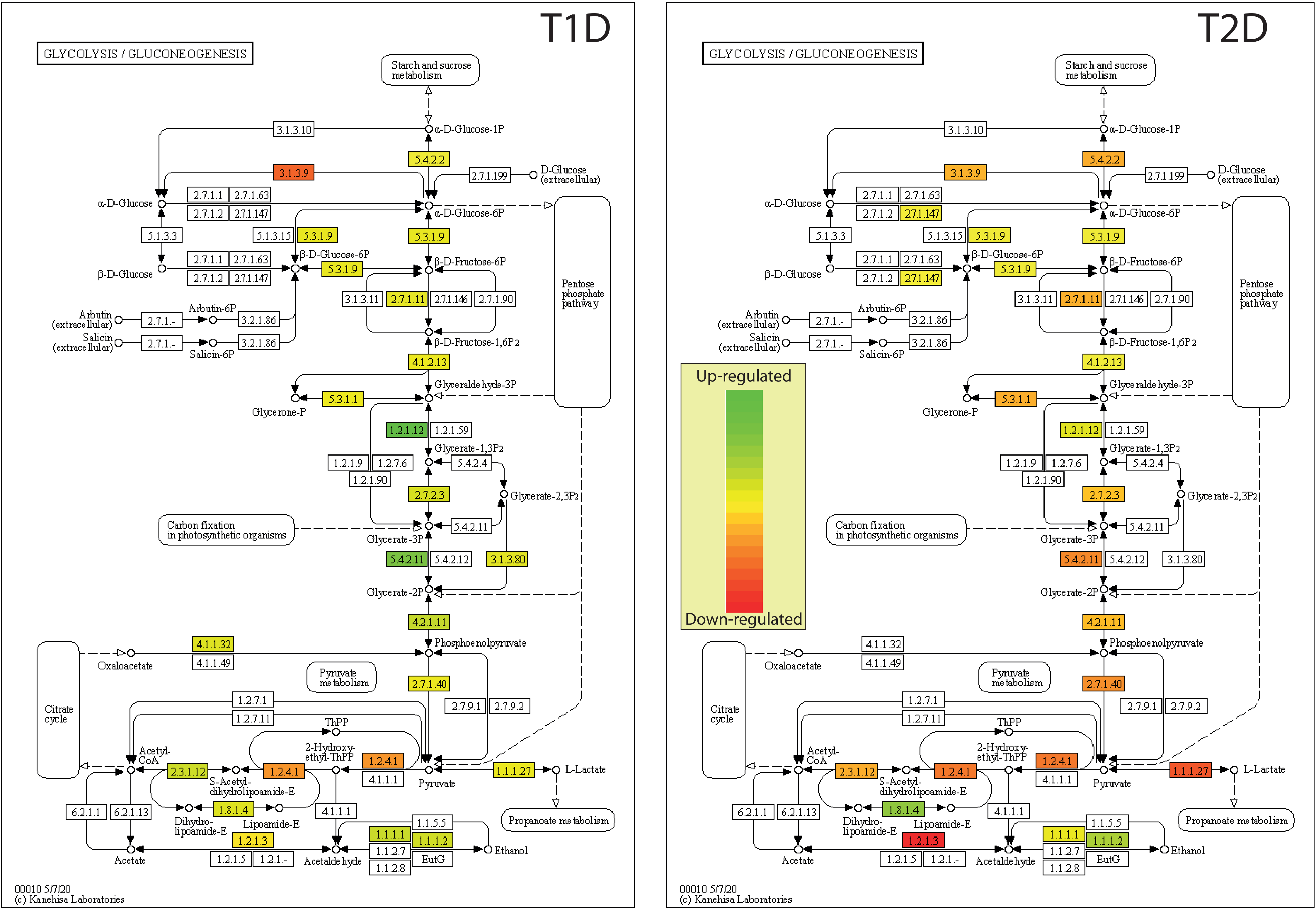

### Supplementary Figure 4

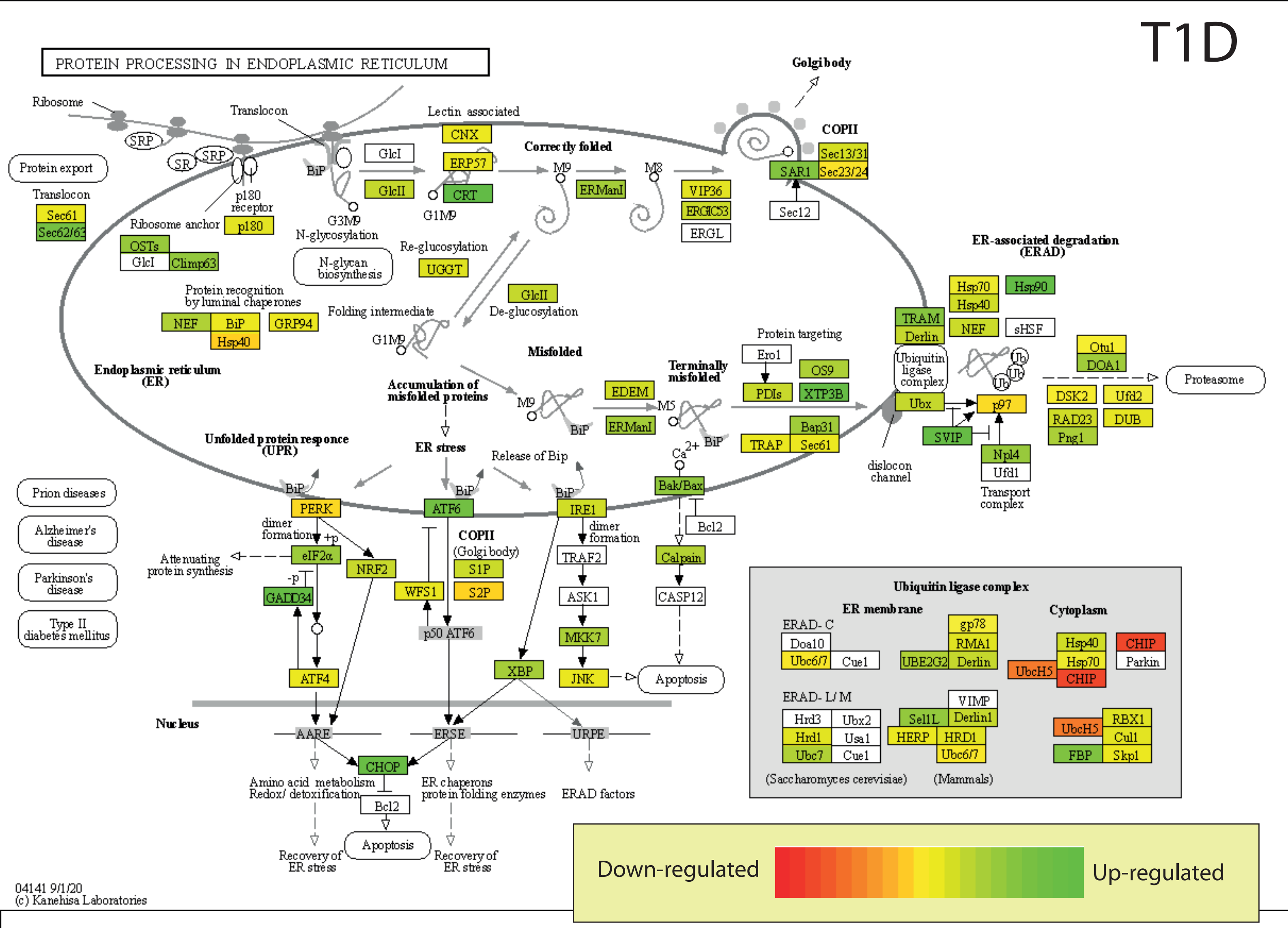

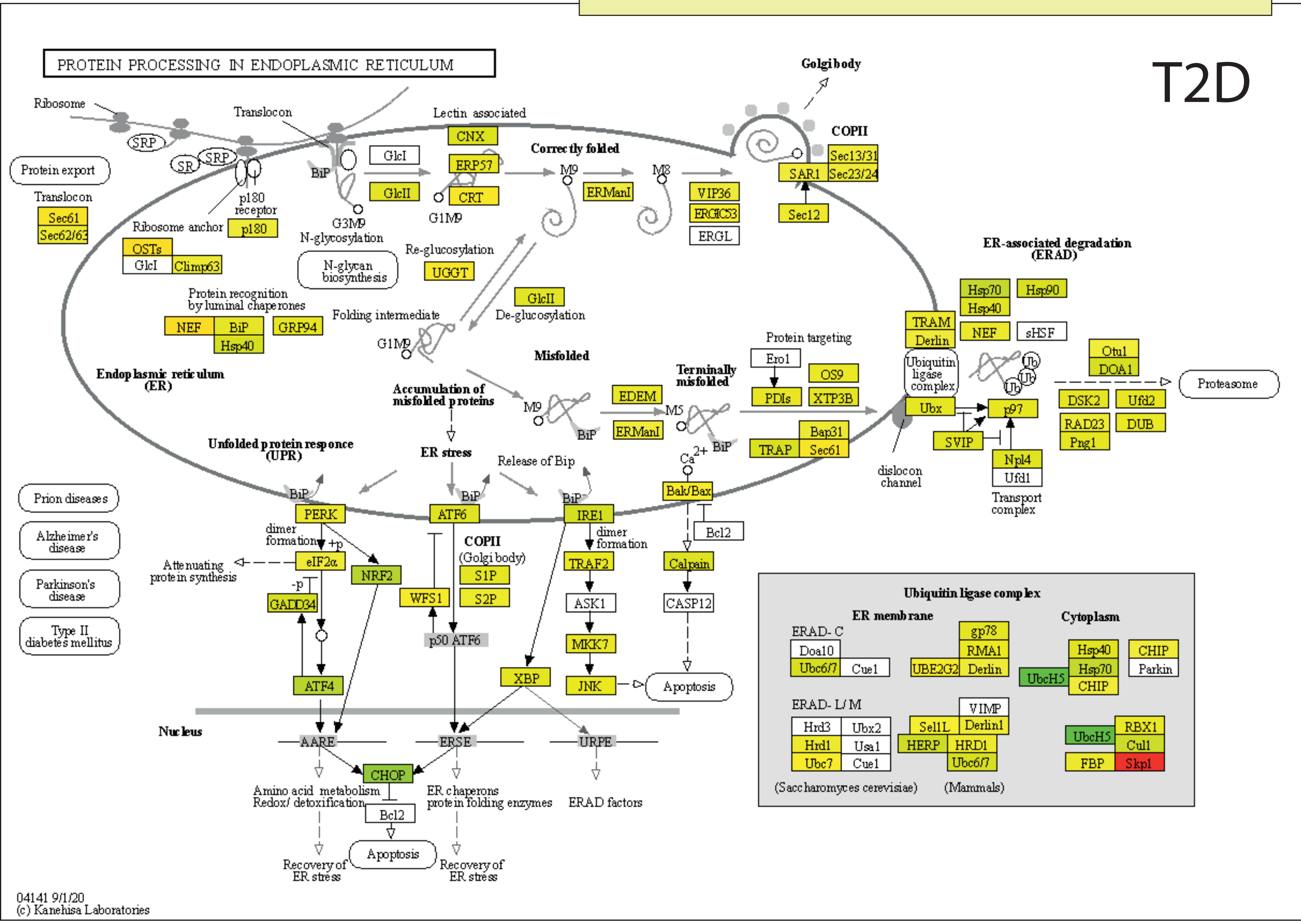
